# Primary human macrophages are polarized towards pro-inflammatory phenotypes in alginate hydrogels

**DOI:** 10.1101/824391

**Authors:** Derfogail Delcassian, Anna A. Malecka, Donaldson Opoku, Virginia Palomares Cabeza, Catherine Merry, Andrew M. Jackson

**Affiliations:** Division of Regenerative Medicine and Cellular Therapies, School of Pharmacy, University of Nottingham, UK; Koch Institute for Integrative Cancer Research, MIT, USA; Host-Tumour Interactions Group, Division of Cancer & Stem Cells, University of Nottingham, UK; Stem Cell Glycobiology Group, Division of Cancer & Stem Cells, University of Nottingham, UK

## Abstract

Dysregulated macrophage function is implicated in a wide range of disorders. *In vitro* hydrogel culture systems are often used as matrices to model and explore the effect of various external stimuli on macrophage polarization and behaviour. Here, we show that 3D alginate hydrogels are not “macrophage inert” and instead help to direct the maturation of primary human macrophages towards specific phenotypes. We compared polarization of M1-like and M2-like cells activated on planar substrates or in 3D alginate hydrogels (with or without adhesion motifs (RGD)). We show that culture in 3D alginate systems selectively alters M2 polarisation following activation; cells show a 2.6-fold increase in CD86 expression compared to cells matured on planar controls, and increase IL1β cytokine secretion even in response to an M2-like stimulus (LPS alone in the absence of IFNγ). Our results suggest that alginate materials may intrinsically stimulate M2 macrophages to acquire a unique polarization state (resembling M2b), characterized by enhanced expression of CD86 and IL1β secretion while retaining low IL12 and high IL10 secretion typical for M2 macrophages. This has important implications for researchers using alginate hydrogels to study macrophage behavior in culture and co-culture systems, as alginate itself may induce direct phenotypic changes independently or in conjunction with other stimuli.

## Introduction

Dysregulated macrophage function is implicated in a wide range of disorders including chronic inflammation^1^, non-healing wounds^2, 3^, and cancer metastasis^4, 5^. Macrophages are traditionally classified on a spectrum from pro-inflammatory (M1) to highly heterogeneous anti-inflammatory (M2), and demonstrate plasticity in transitioning between phenotypes.^5–11^ In the tumor microenvironment, the presence or absence of specific macrophage phenotypes is thought to play an important role in disease progression and response to therapeutics. The pro- or anti-inflammatory nature of the tumor microenvironment is driven, in part, by the interaction of tumor associated macrophages (TAMs) with cancer cells.^4, 10, 12, 13^ There is particular interest in understanding how cancer cells can modulate the phenotype and behavior of these immune cells, and so regulate the tumor microenvironment. Many *in vitro* model systems therefore seek to understand the behavior of macrophages and their interaction with the extracellular matrix and other cell types.

Hydrogels are widely used as model *in vitro* systems to study cellular interactions occurring in a variety of microenvironments.^14–17^ These culture systems can be fabricated from a broad range of polymer sources, including both synthetic and natural materials.^18–22^ Alginate is a natural polysaccharide polymer derived from seaweed, which consists of long chains of guluronic acid (G) and mannuronic acid (M) typically arranged as a block co-polymer.^20^ Alginate can be reversibly crosslinked *via* cationic interactions with divalent cations, and various ligands can be crosslinked into the polymer backbone.^15, 20, 23^ Following reversible hydrogel dissociation, cells can be recovered for further analysis, making alginate a particularly useful platform system to model and probe macrophage behavior in co-culture systems *in vitro.*

There is emerging evidence that modifications to alginate may affect macrophage cell behavior and phenotype.^24^ Tuning alginate stiffness, chemical backbone composition, or the functional groups attached to alginate can impact the phenotype of mouse macrophage cell lines cultured on alginate *in vitro.*^24–26^ Although precise structure function relationships have yet to be identified, it appears that functionalized alginate is not a “macrophage-inert” medium. As alginate hydrogels are widely used in *in vitro* models to study macrophage behavior, we sought to understand whether alginate itself affects primary human macrophage polarization *in vitro*.

Here, we use 3D culture systems to explore the effect of alginate and alginate functionalized with RGD adhesion ligands on primary human macrophage polarization. To our knowledge, this is the first study on the effect of 3D alginate culture on primary human macrophage polarization *in vitro*. We show that human macrophage polarization is altered in 3D alginate culture systems. This has important implications for researchers using alginate hydrogels to study macrophage behavior in culture and co-culture systems, such as those used to model the tumor microenvironment, as alginate itself may selectively induce specific functions of macrophages independently or in conjunction with other stimuli therefore directing macrophage polarization.

### Macrophage polarization in 3D alginate hydrogel systems

We compared the polarization of primary human macrophages isolated from peripheral blood monocytes *in vitro* using traditional well plates (planar culture) or 3D alginate hydrogel culture (3D culture) (Figure 1). This allowed us to model the response of non-tissue specific macrophages. Various chemical stimuli can be used to direct macrophages towards a specific phenotype on a spectrum from pro-inflammatory M1-like behavior to anti-inflammatory M2-like behavior.^5^ Here, we used GMCSF to generate immature M1-like, or MCSF for immature M2-like cells. Then, immature M1 and M2 cells were isolated and cultured on tissue culture plastic or in 3D alginate hydrogel systems, and activated for 48 hours with LPS and IFNγ for a mature M1-like phenotype, and LPS alone for a mature M2-like phenotype (Figure 1).

**Figure 1:**
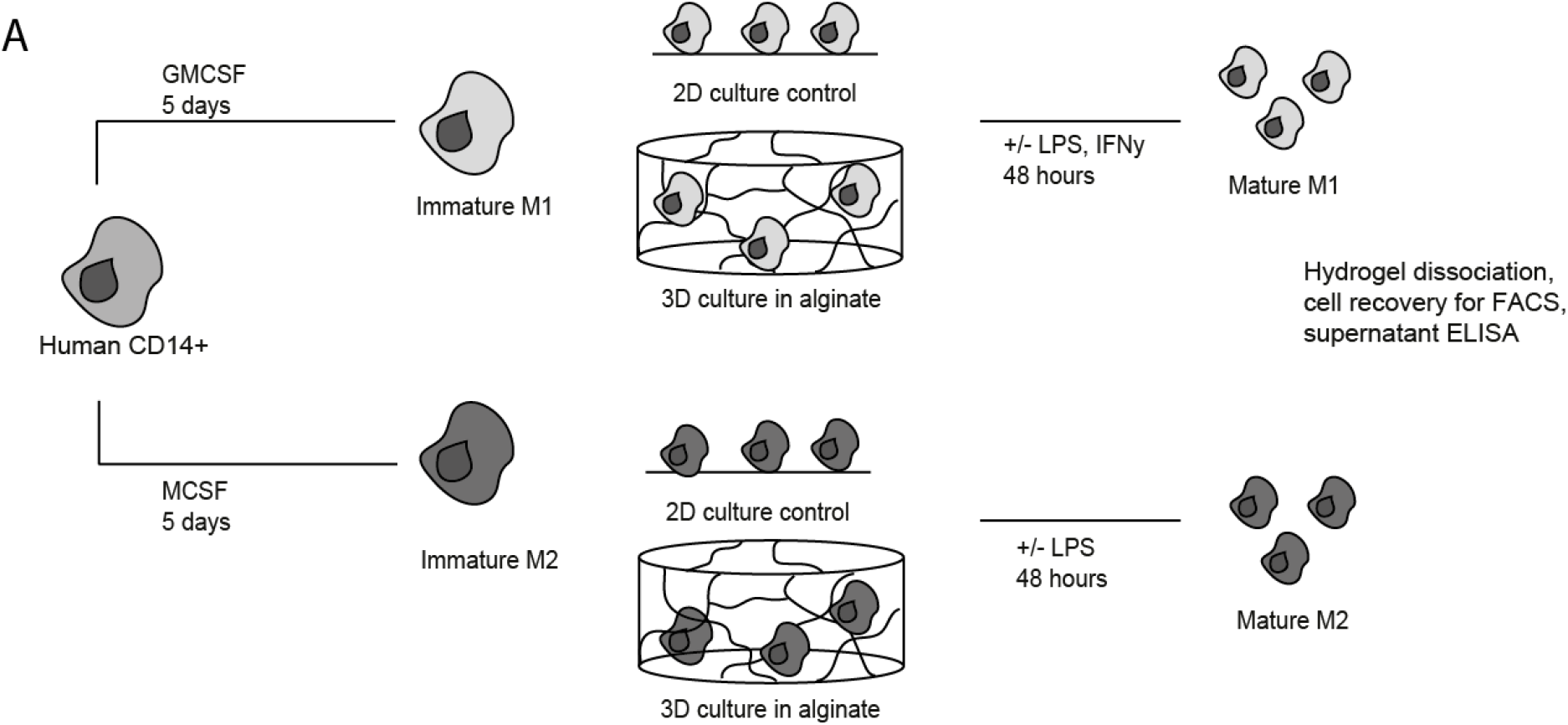
*In vitro* 3D polarization of human macrophages. Primary human CD14+ cells were isolated from peripheral human blood, and cells were cultured to generate immature M1 or M2 cells. Immature cells were then encapsulated within alginate in a 3D culture system, or were treated on planar polystyrene controls, and polarized towards a mature M1-like or M2-like phenotype. After 48 hours of stimulation, supernatants were harvested and cells were dissociated from planar surfaces or hydrogel materials for flow cytometry analysis.

To analyse cell phenotype and cell function *in vitro*, supernatants were collected after 48 hours, gels were dissolved, and cells were isolated for flow cytometry analysis. We evaluated cell surface marker expression to simply classify the *in vitro* polarization of primary human macrophages in planar and 3D culture. Figure 2 shows expression levels of typical M1 markers CD86 and MHC II on macrophages treated in these conditions. As expected, in planar culture systems M1 cells increased expression of MHC II (Figure 2A1) and CD86 (Figure 2C1) following stimulation, whilst M2 cells showed little change in CD86 on maturation (Figure 2B1), and a slight increase in MHC II expression (Figure 2).

**Figure 2:**
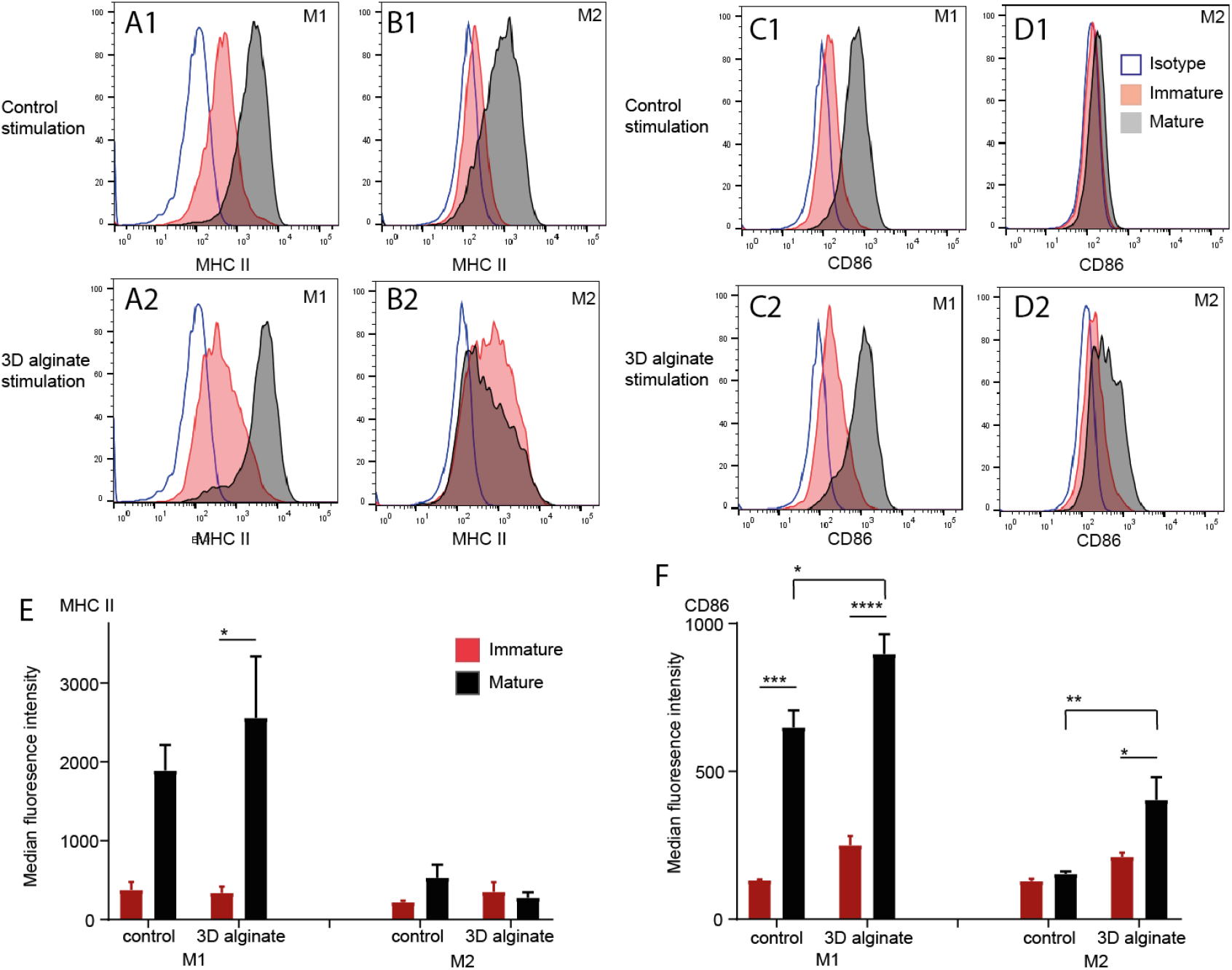
Cell surface marker expression following activation. Flow cytometric analysis of surface expression of A,B) MHC II and C,D) CD86 following maturation. Cells were cultured in (A-Di) planar or (A-Dii) 3D alginate culture systems. Cells were gated as live, single cells and representative data from one donor compared to an immature M1 or M2 isotype control is shown. A-Diii) Average median fluorescence intensity of MHC II and CD86 cell surface expression from three independent human donors. Graphs show the average of 3 different human donors and error bars represent standard error in the mean. Statistics were performed using GraphPad Prism, using two-way ANOVA and Turkey’s multiple comparison tests to analyse differences between immature and mature cells in planar and 3D systems, and between mature cells in these conditions. *p<0.05, **p<0.01, ***p<0.001, ****p<0.0001.

In 3D systems, M1 cells significantly increased expression of MHC II on maturation (as expected), and there was no significant change in MHC II expression in 3D culture conditions compared to planar controls (Figure 2A2). In contrast, CD86 expression was significantly increased in both M1 and M2 cells matured in 3D alginate structures compared to planar controls (Figure 2F; 1.4-fold (M1) and 2.6-fold (M2) increase in CD86 expression). Cells showed increased basal CD86 expression compared to planar controls even pre-activation (Figure 2F). This increase was particularly important for M2-like cells, whose CD86 expression is significantly upregulated 2.6-fold to display an “intermediate” level of expression between typical M1-like and M2-like cells cultured on planar controls. This suggests that stimulation in our 3D culture system leads to an increase in CD86 expression in both M1-like and M2-like cells, resulting in altered activation compared to planar controls. Next, we explored whether this upregulation in CD86 expression was altered in the presence of adhesion ligands.

### Macrophage polarization in 3D alginate hydrogel systems presenting adhesion ligands

Immature macrophages cultured in 3D showed increased basal expression of CD86. Because the co-stimulatory molecule CD86 serves to strengthen the association between macrophages and T-cells,^27, 28^ we hypothesized that immature macrophages upregulate CD86 in 3D alginate matrices to increase their association with the surrounding matrix. To test this, we generated three alginate hydrogels with matched rheological properties and increasing concentrations of an RGD motif. RGD is a tri-peptide adhesion motif which is found in extracellular matrix proteins (e.g. fibronectin and collagen) and binds to cell surface adhesion receptors such as integrins.

To maintain similar mechanical properties between conditions as the concentration of RGD in the alginate phase was increased, we made gels with 0mg/ml to 6.5mg/ml RGD using co-crosslinked RGD conjugated alginate and unfunctionalised alginate. We compared Young’s Modulus to characterize the mechanical properties of these gels using parallel plate rheometry at increasing strain (Figure 3A-C). The hydrogels were viscoelastic, with a storage modulus at least one order of magnitude higher than the loss modulus, and presented Young’s Moduli between 8-12kPa. There was no significant difference in Young’s Modulus between the three conditions. However, gels with higher concentrations of RGD-alginate exhibited slightly higher Young’s Moduli compared to unfunctionalised controls.

**Figure 3:**
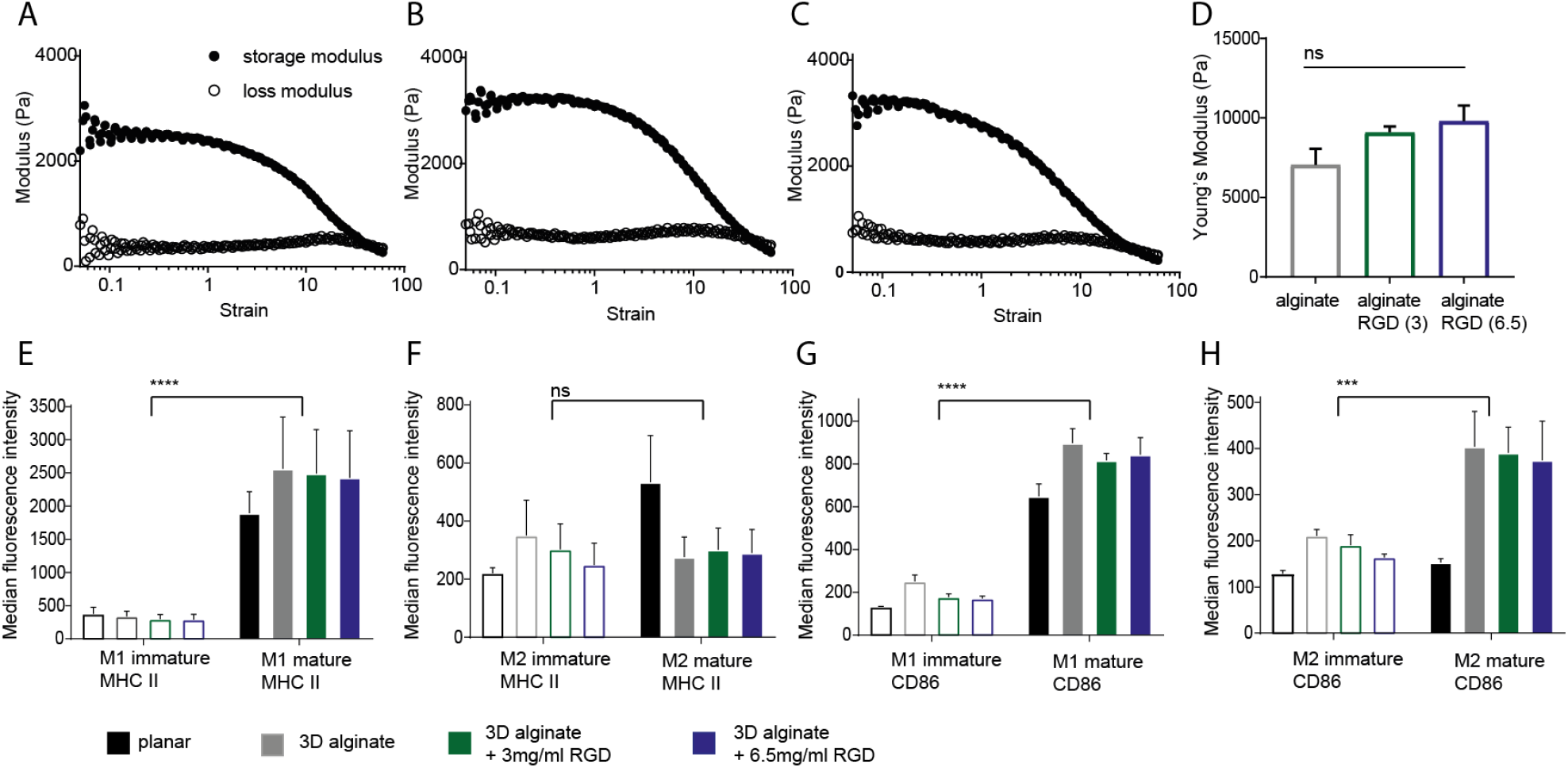
RGD alginate interfaces for 3D macrophage culture. Alginate materials containing 0mg/ml, 3mg/ml or 6.5mg/ml alginate-RGD were fabricated and cross linked in calcium chloride. A-C) Parallel plate rheometry was used to measure storage and loss modulus of the materials at constant amplitude and increasing strain. Graphs show moduli averaged from three different gels of each kind. D) Young’s Modulus was calculated for each gel using E=2G (1+*ν*) where *ν* is Poisson’s ratio (~0.5 for these equations) at 0.1% strain, by averaging values from at least three different gels. E-H) Flow cytometric analysis of surface expression of E, F) MHC II and G, H) CD86 following maturation. Cells were cultured in planar or 3D alginate culture systems. Cells were gated as live, single cells and average median fluorescence intensity of MHC II and CD86 cell surface expression from three independent human donors is shown, error bars represent standard error in the mean. Experiments were performed independently for at least three biological donors. Statistics were performed using GraphPad Prism, using Turkey’s multiple comparison tests to analyse differences between immature and mature cells. *p<0.05, **p<0.01, ***p<0.001, ****p<0.0001.

We examined whether the presence of RGD altered macrophage phenotype. For immature M1 and M2 cells, increased RGD concentration resulted in a slight but non-significant decrease in MHC II and CD86 and this was more notable on the M2 subset. On maturation, cells within RGD-alginate hydrogels displayed similar MHC II and CD86 expression levels to cells cultured in alginate hydrogels. As noted for immature cells, surface expression of CD86 in M2 cells altered in the presence of RGD; cells matured in gels functionalized with increasing concentrations of RGD showed slightly decreased CD86 expression compared to cells cultured in an unfunctionalised alginate, again, this was not significant.

Interestingly, cells cultured in hydrogels expressed similar levels of CD86 markers, regardless of the RGD concentration. This suggests that the alginate matrix itself (rather than the availability of adhesion ligands such as RGD) modulates the altered expression of MHCII and increased expression of CD86, particularly in the M2-like phenotype.

### Cytokine secretion following stimulation

We analysed cytokine expression in cell culture supernatants following stimulation (Figure 4) to further characterize the impact of planar or 3D alginate culture on macrophage behaviour. As expected, cells cultured in the planar systems showed significantly elevated IL12, TNFα and IL1β secretion in M1 polarised cells compared to M2 cells (Figure 4A,C), whilst M2 cells demonstrated significantly increased IL10 production (Figure 4B). In 3D culture systems, cells showed similar trends for IL12, TNFα and IL10 production (Figure 4E-G) however cytokine levels present in cell supernatants were significantly reduced. Interestingly, IL1β secretion was elevated in M2 cells matured in 3D culture systems (Figure 4H). Next, we compared the ratio of cytokine expression in M1 cells over M2 cells for cytokines IL12 and IL1β (Figure 4J). This method minimizes any variations in absolute cytokine levels in supernatants due to cytokines with different charges or sizes being retained differently within a hydrogel, or due to delayed cytokine release kinetics from cells encapsulated within 3D hydrogels, and instead compares the ratio of secretion between M1 and M2 cells in the same maturation matrix.

**Figure 4:**
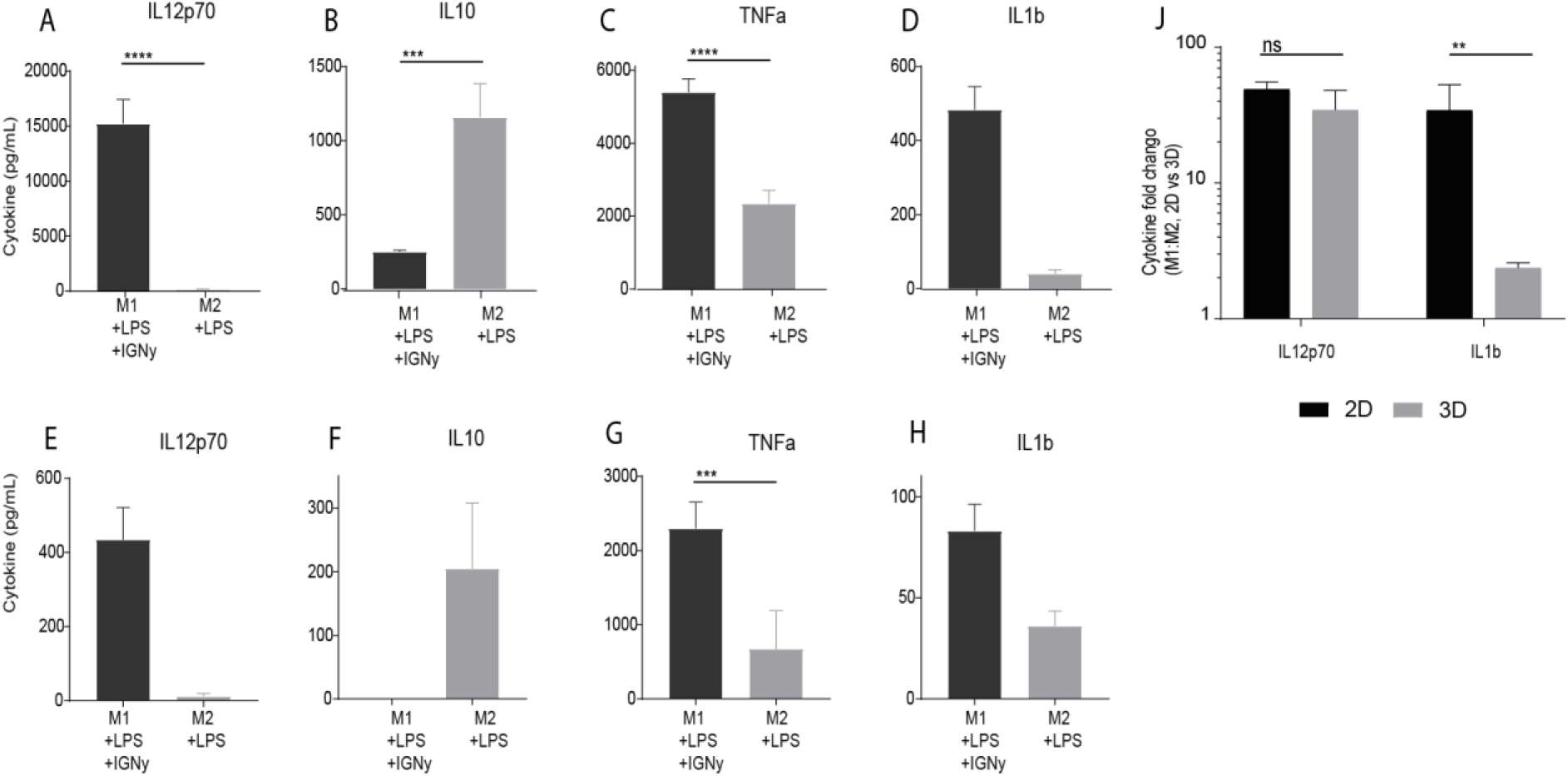
Cytokine secretion from planar and 3D macrophage cultures. A-H) Macrophages were polarized in planar (A-D) or 3D (E-H) systems, and supernatants were analysed after 48 hours of stimulation. Supernatants were tested for IL12, IL10, TNFa and IL1β levels. J) The fold change in cytokine secretion was calculated for matched M1:M2 cells, and compared between planar and 3D systems. Average cytokine expression from 3 donors is plotted, error bars represent standard error in the mean. Statistics were performed using GraphPad Prism, using Turkey’s multiple comparison tests to analyse differences between M1 and M2 cells, and a Mann Whitney T-test to analyse fold changes. *p<0.05, **p<0.01, ***p<0.001, ****p<0.0001.

We found that the ratio of IL12 production between M1 and M2 cells was consistent in both planar and 3D culture, averaging 35-50 fold higher secretion in M1 cells compared to M2. In contrast, IL1β cytokine secretion was significantly altered between planar and 3D systems. Following maturation in planar systems, M1 cells produce around 30 fold more IL1β than M2 cells. However, in 3D alginate systems, M1 cells produce only 2-3 fold more IL1β (Figure 4J). This supports our earlier CD86 and HLA-DR expression data and suggests that primary human macrophage polarization in 3D alginate hydrogel systems is selectively modified by alginates, especially for M2-like cells. Upon maturation in alginate hydrogels, M2 cells increase CD86 expression and increase secretion of IL1β, in contrast to cells cultured on tissue culture plastic controls.

The mechanism driving M2-like cells to secrete IL1β in response to these alginate hydrogels has not yet been fully elucidated. Several sub-sets of M1- and M2-like cells have been described (including M2a, M2b, M2c and M2d) and macrophages appear able to dynamically transition between these phenotypes.^5, 12, 29^ Here, both the increase in CD86 expression and IL1β secretion are seen following additional maturation with LPS, suggesting that alginate provides an additional stimuli to these cells that is absent from planar culture systems. High CD86 expression and IL1β secretion together with high IL10 and low IL12 secretion are markers of M2b macrophages which can be generated in the presence of LPS and immune complexes.^30, 31^ The increase in CD86 and IL1β expression without an effect on the IL12/IL10 ratio suggest selective action of the alginates. Therefore it may be hypothesized that alginates act in similar way to immune complexes. Furthermore, IL1β is a pro-inflammatory cytokine often upregulated in response to stimulation *via* Toll Like Receptors (TLRs).^32^ Several TLR agonists have been derived from short-chain polysaccharide systems,^32, 33^ and it is possible that alginate materials may therefore intrinsically stimulate IL1β secretion in primary human macrophages *via* this pathway.

## Conclusions

We show that macrophage maturation in alginate materials alters macrophage polarization through selective increase in specific pro-inflammatory markers. We analysed cell surface marker expression (CD86, MHCII) and cytokine secretion profiles, and demonstrate that activation within alginate gels alters M2 polarisation compared to planar controls. Although M1 cells mature as expected, M2 cells significantly increase CD86 expression (2.6 fold) and IL1β cytokine secretion upon maturation in 3D cultures compared to planar tissue culture plastic controls. At the same time other typical pro-inflammatory markers such as IL12, TNFa and HLA-DR as well as anti-inflammatory IL10 remained unchanged. The inclusion of adhesion motifs (RGD) did not significantly alter cell polarization for either the M1 or M2 phenotype. These findings demonstrate the specificity of alginate action and further highlight the requirement to assess a wide array of markers when investigating the immunogenicity of biomaterials.

To our knowledge, this is the first demonstration that primary human macrophage polarization, especially of M2-like cells, is altered in 3D alginate systems. This has important implications for researchers using alginate hydrogels to study macrophage behavior in co-culture systems or as implantable alginate biomaterials. Model systems which employ alginate as a polymer matrix for cell culture and co-culture studies (such as those used to model cell behavior in the tumor microenvironment) may inadvertently polarize or prime macrophages towards specific phenotypes. CD86, IL1β and IL10 are typical markers of a specific immunoregulatory M2b subtype of macrophages, which play an important role in cancer progression and development. ^34–36^ Their polarization can normally be achieved by activation of human macrophages with LPS in the presence of immune complexes; alginate materials may therefore intrinsically stimulate primary human macrophages *via* this pathway. Careful consideration should therefore be given to the design of these alginate systems, especially when investigating anti-inflammatory macrophage function.

## Materials and methods

### Reagents

All reagents were endotoxin-free. Recombinant human GM-CSF was purchased from Peprotech, and IFN-γ from R&D Systems. Ultrapure TLR4-agonist (*Salmonella Minnesota* LPS) was purchased from InvivoGen. Recombinant human M-CSF was obtained from ImmunoTools. Antibodies and matched isotype controls were obtained from Miltenyi.

### Cell isolation and generation of monocyte-derived macrophages

Human macrophages were generated from CD14^+^ monocytes, isolated from healthy donors (fresh blood or Buffy Coats), from the National Blood Transfusion Service in accordance with the approval of the relevant ethical review boards. Samples were fractioned to isolate PBMCs using a gradient centrifugation using Histopaque 1.077 (Sigma Aldrich), and CD14+ cells were isolated using anti-CD14 magnetic microbeads (Miltenyi Biotech). Monocytes were cultured in RPMI-1640 medium supplemented with 10% FCS and 1% sodium pyruvate (all from Sigma) in ultra-low attachment 6-well plates. To polarise macrophages, recombinant human GM-CSF (20U/mL) or M-CSF (10ng/mL) was added and the cells were cultured for 6 days, adding additional supplemented medium on day 4. Macrophages were obtained on day 6, and their quality and purity assessed by flow cytometry.

### Hydrogel synthesis and analysis

We used sterile sodium alginate crosslinked with divalent calcium ions as our alginate hydrogel. Sterile sodium alginate (G/M ≥1.5, endotoxins ≤ 100EU/g, Novamatrix) was dissolved in saline or complete media (RPMI-1640 Medium (Sigma), supplemented with 10% DC-Fetal Bovine Serum (FBS) protein, 1% streptomycin, 1% penicillin, 1% sodium pyruvate) at concentrations between 1-3% overnight on a roller at 37°C. Cells were encapsulated at 0.5×10^6^ cells/mL, and added to the alginate before cross-linking. 400uL of hydrogel was cast in each well of a 48 well plate, and cross-linked with 0.1 M CaCl_2_ for 45 minutes at room temperature, before excess crosslinking solution was removed and wells washed with PBS. Parallel plate rheometry was used to determine Youngs Modulus, using the equation *E*=2G (1+*ν*), where E is Young’s Modulus, G is Modulus at a given strain/frequency, and *ν* is Poisson’s ratio (Figure 3D).

### Stimulation of macrophages

Immature Mφ were counted, washed and cultured on tissue culture plastic or in 3D alginate hydrogel systems. Mφ were activated for 48 hours withLPS (500ng/mL) and IFNγ (1000 U/mL) to direct cells towards a mature M1-like phenotype, and LPS (500ng/mL) alone for a mature M2-like phenotype. To analyse cell phenotype and cell function *in vitro*, supernatants were collected after 48 hours, gels were dissolved, and cells were isolated for flow cytometry analysis. Supernatants were stored at −20°C before ELISA analysis.

### Characterisation of phenotype by flow cytometry

Macrophages from planar cultures were detached by gentle agitation in PBS. Macrophages in gels were collected after dissolving gels in 150mM EDTA for 15min at 37°C. Cells were washed twice in PBS and stained in FACS buffer (PBS, 1% FBS, 0.1% sodium azide) with CD86 (clone FM95) and HLA-DR (clone AC122) or matched isotypes control antibodies (all Miltenyi). Samples were fixed in 2% formaldehyde (Sigma) and stored in the dark at 4°C until acquisition. Samples were acquired on MACSQuant Flow Cytometer and analysed using FlowJo software (Treestar Inc.).

### Determination of cytokine secretion

Diluted cell culture supernatants were assayed by ELISA for IL-12 (BD Bioscience, UK) and IL-10, IL-1β and TNFα (R&D Systems, UK) according to the manufacturer’s instructions. Assays did not significantly cross-react with other proteins (sensitivities were 7.8, 15pg/mL, 3.9 and 15.6pg/mL respectively).

### Statistical analysis

The statistical analysis was performed using GraphPad Prism software. Turkey’s multiple comparison was used to analyse differences between individual conditions for immature and mature cells, and two-way ANOVA to analyse differences between groups. P-values of less than 0.05 (p-value < 0.05) were considered significant, and p values are expressed as *p<0.05, **p<0.01, ***p<0.001, ****p<0.0001.

## Acknowledgements

DD acknowledges funding from an EPSRC E-TERM Fellowship which helped to support this work. AM was funded by a University of Nottingham Vice Chancellor’s Research Excellence Scholarship.

